# Retrospective image analysis for long-term demography using Google Earth imagery

**DOI:** 10.1101/2025.09.02.673673

**Authors:** Erola Fenollosa, Garrett Cohen, Roberto Salguero-Gómez

## Abstract

1. Ecosystems are rapidly degrading. Widely used approaches to monitor ecosystems to manage them effectively are both expensive and time consuming. The recent proliferation of publicly available imagery from satellites, Google Earth, and citizen-science platforms holds the promise to revolutionising ecological monitoring and optimising their efficiency. However, the potential of these platforms to detect species and track their population dynamics remains under-explored.
2. We introduce a fast, inexpensive method for retrospective image analysis combining current ground-truth data with historical RGB imagery from Google Earth to extract long-term demographic data. We apply this method to three case studies involving two major Mediterranean invasive plant taxa with contrasting growth forms. Specifically, we: (1) utilise deep learning to automatically detect individuals of prickly pear (*Opuntia* sp.) across various Mediterranean habitats and image resolutions; (2) reconstruct 10 years of spatially explicit recruitment rates for *Opuntia* along a climatic gradient; and (3) quantify nearly 20 years of growth dynamics for the clonal invader *Carpobrotus* sp. in two contrasting environments.
3. Our object detection model, trained with Google Earth imagery, achieves 60-80% success in identifying individuals of *Opuntia*, regardless of habitat type. Model performance increases with target species colour consistency and contrast, as well with the usage of basic data augmentation techniques. Detection is constrained by individual area (<4 m²) but captures 80% of the examined population.
4. Beyond detection, our time-series analysis of publicly available imagery enables detailed population monitoring. With 10-year image series available for Spain, Greece, and the UK, and 20 years for Portugal, we successfully estimate annual recruitment and growth rates and their climatic sensitivity, identify productive and unproductive years, estimate individual age, characterise population structure, model size-age relationships, and identify recruitment hotspots for targeted management.
5. Our pipeline opens new avenues for cost-effective, large-scale demographic monitoring by retrospectively harnessing open-access imagery. While demonstrated here with invasive plants, we discuss the broad applicability of our approach across taxa and ecosystems. The use of retrospective image analysis for long-term demography with Google Earth imagery has the potential to expedite conservation decisions, support effective restoration, and enable robust ecological forecasting in the Anthropocene.

## 1. Introduction

Monitoring natural populations is key to understanding and managing biodiversity (Lindenmayer and Likens, 2010). However, traditional methods—such as repeated field observations or individual tracking—are often time-consuming, error-prone, and resource-intensive (Lindenmayer et al. 2012; Gamelon et al. 2021; Besson et al. 2022). Consequently, population studies are frequently limited to small spatial scales or short timeframes, hindering ecological inference and delaying interventions (Salguero-Gómez et al. 2014; 2016; Compagnoni et al. 2021; Römer et al. 2023). Yet long-term data are essential to quantify key demographic processes such as survival, recruitment, and growth (Blanc and Thrall 2024; Stroud and Ratcliff 2025). Moreover, transient (*i.e.*, short-term) dynamics (Stott et al. 2011) and environmental stochasticity render the forecasting of long-term populations even more complicated (Caswell 2001; Boyce et al. 2006; Hock et al. 2024).

Extensive populations mapping and monitoring reveals large-scale ecological patterns, including shifts in species distributions, population densities, or transitions between vegetation types (Tingley and Beissinger, 2009; Lenoir and Svenning, 2014; Pisa et al. 2019; Steel et al. 2021). Such analyses can help identify drivers of ecological change—like climate shifts or invasive species—and support targeted conservation actions (Bürgi et al. 2005; Edmunds, 2021; Buchan et al. 2022). Localising individual-level recruitment and mortality from images has been proven useful to detect disturbance responses (Jaime et al. 2023) and can contribute flagging early warning signs of ecosystem degradation or recovery (Visser et al. 2014; Pasquarella et al. 2016; Alibakhshi 2023; Poulton et al. 2024).

Recent advances in high-resolution imagery and machine learning offer powerful tools for ecological monitoring (Lahoz-Monfort and Magrath 2021; Besson et al. 2022; Cavender-Bares et al. 2022; Reynolds et al. 2025). A growing body of work now integrates diverse image sources—ranging from online photos to satellite imagery—to study ecological dynamics (Leighton et al. 2016; Depauw et al. 2022; Pasquarella et al. 2016). Aerial imagery is already used to track species from elephants to herbs (Duporge et al. 2020; Young et al. 2022; Zagajewski et al. 2024), often paired with deep learning algorithms to automate detection (Wäldchen and Mäder 2018; Retallack et al. 2022).

Google Earth Pro provides public high-resolution imagery for ∼98% of terrestrial surfaces (Yu and Gong 2011; Liang et al. 2018; Google 2019; Yang et al. 2022), including historical imagery often finer than that of open satellite sources (15 cm vs. 30 cm–25 m/pixel; EOS Analytics 2025). Despite its availability, ecological applications of Google Earth remain scarce. Some studies have mapped vegetation change or species presence—e.g., palms (Minnich et al. 2011), bamboo forests (Watanabe et al. 2020), and invasive trees (Visser et al. 2014)—but typically limited to binary comparisons across two time points. No existing study has systematically tracked individual-based population dynamics using Google Earth across multiple years.

The monitoring of invasive species is key to designing targeted management actions to protect biodiversity (Singh et al. 2024). Invasive species are one of the main threats to global biodiversity (Roy et al. 2024; Schwindt et al. 2024), with global costs estimated around €144.8 billion/year (Diagne et al. 2021). Classical monitoring methods are often costly and ineffective (Javed et al. 2025), emphasising the critical need for timely monitoring and early intervention (Ahmed et al. 2022). Novel monitoring technologies offer the potential to revolutionise invasive species management by reducing both detection costs and response times (Zaka and Samat 2024; Javed et al. 2025). Invasive plants are ideal candidates for image-based tracking, as they are sessile, fast-growing, and visually conspicuous (Visser et al. 2014).

Here, we present a novel pipeline for reconstructing long-term demographic data using retrospective, open-access Google Earth imagery. We demonstrate its application with two morphologically distinct invasive plant taxa, combining field-based ground truthing with deep learning detection. Analysing 20 years of imagery across seven sites in four European countries, we reveal sustained recruitment dynamics and climate-driven responses. Finally, we identify conditions under which our approach is most effective, offering a blueprint for broader applications in biodiversity monitoring and management.

## 2. Material and Methods

Our pipeline builds on the idea that confirming species persistence over time is easier than detecting them initially (Bewley et al. 2016; James and Bradshaw, 2020). The open access of Google Earth imagery enables the reconstruction of species’ spatiotemporal dynamics after field-based geopositioning (**Figure 1**).

**Figure 1.**
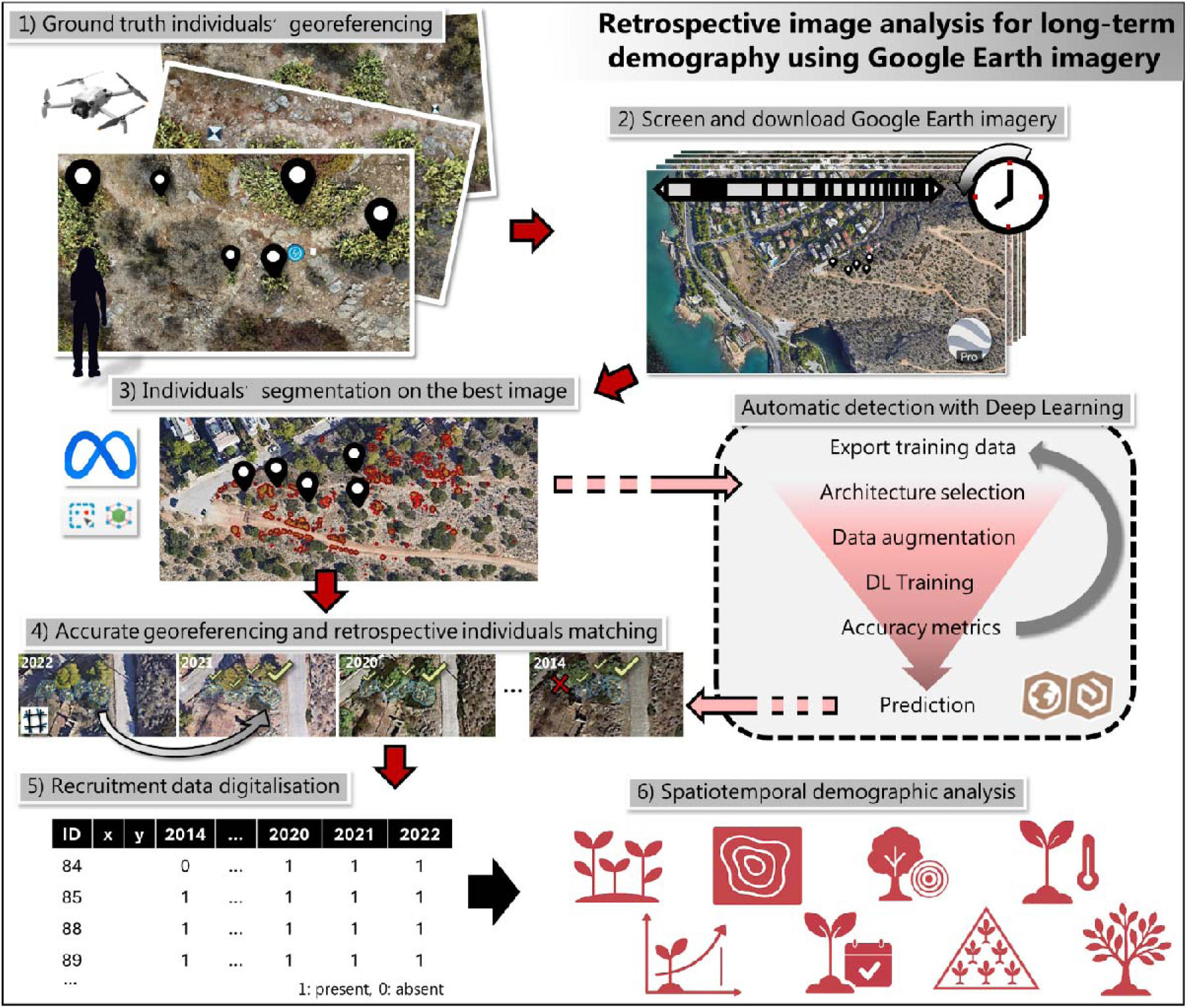
Pipeline for retrospective image analysis for long-term population monitoring using Google Earth imagery. With only a field visit to acquire ground truth data of individual’s presence (Step 1), it is now possible to acquire data of the spatiotemporal population dynamics of invasive species. Step 1 is crucial to determine the current presence of the individuals that will be tracked back in time. Both GPS positioning and UAV flights that provide high quality imagery can be used. After delimiting the specific position of individuals in the population, we can proceed to download Google Earth Pro Imagery (Step 2), screening all possible images across time. Combining information regarding an individual’s position and Google Earth imagery, Step 3 involves segmenting the extension of all the individuals, which can be done using AI-assisted segmentation. At this point, we can use the segmented polygons to train deep learning algorithms and detect further individuals. All images need to be accurately georeferenced to match individuals back in time (Step 4). Once all images have been georeferenced, we can acquire recruitment data by confirming the presence of individuals on the multiple time steps and potentially include their size (Step 5). Recruitment data can be used for multiple purposes, including full population size and structure estimates, quantifying annual recruitment rates, and exploring their climatic sensitivity, estimating individual age, modelling age-size relationships, quantifying annual growth, building individual-based models and identifying high and low productivity areas (Step 6). Video-tutorials and a standardised protocol for this analytical pipeline can be found in **Supplementary Material 1**, as well as this YouTube playlist: https://www.youtube.com/playlist?list=PL_LKE-yTi9kBXfw_qDdJCQ3Sxu2fjGvDD

### Species and study sites

We focused on two major invasive plant taxa in Mediterranean ecosystems: *Opuntia* spp. (e.g., *O. ficus-indica*, *O. stricta*, *O. dillenii*, *O. amyclaea*) and *Carpobrotus* spp. (*C. edulis*, *C. acinaciformis*) (Erre et al. 2009; Campoy et al. 2018; Novoa et al. 2019). *Opuntia* is a tall, upright cactus with fleshy, spiny cladodes; *Carpobrotus* forms dense ground-cover mats with rooting trailing stems, often displacing native vegetation (Bogdan et al. 2021; Fenollosa et al. 2025a).

We surveyed five *Opuntia* and two *Carpobrotus* populations across their Mediterranean and latitudinal ranges, respectively (**Figure 2**). At each site (0.2–2 ha), we geolocated >100 individuals and captured high-resolution RGB imagery using a DJI Mini 4 Pro (0.436 cm/pixel GSD). Flights were planned in PixPro (Pixpro Ltd, UK) with 85% overlap, 10 m altitude, 2 s intervals, and a -90° camera angle to optimise stitching in Pix4DMapper (Pix4D SA, Switzerland). We used the GPS position of each individual plant and drone imagery as ground truth data of individual locations to train deep learning object detection models.

**Figure 2.**
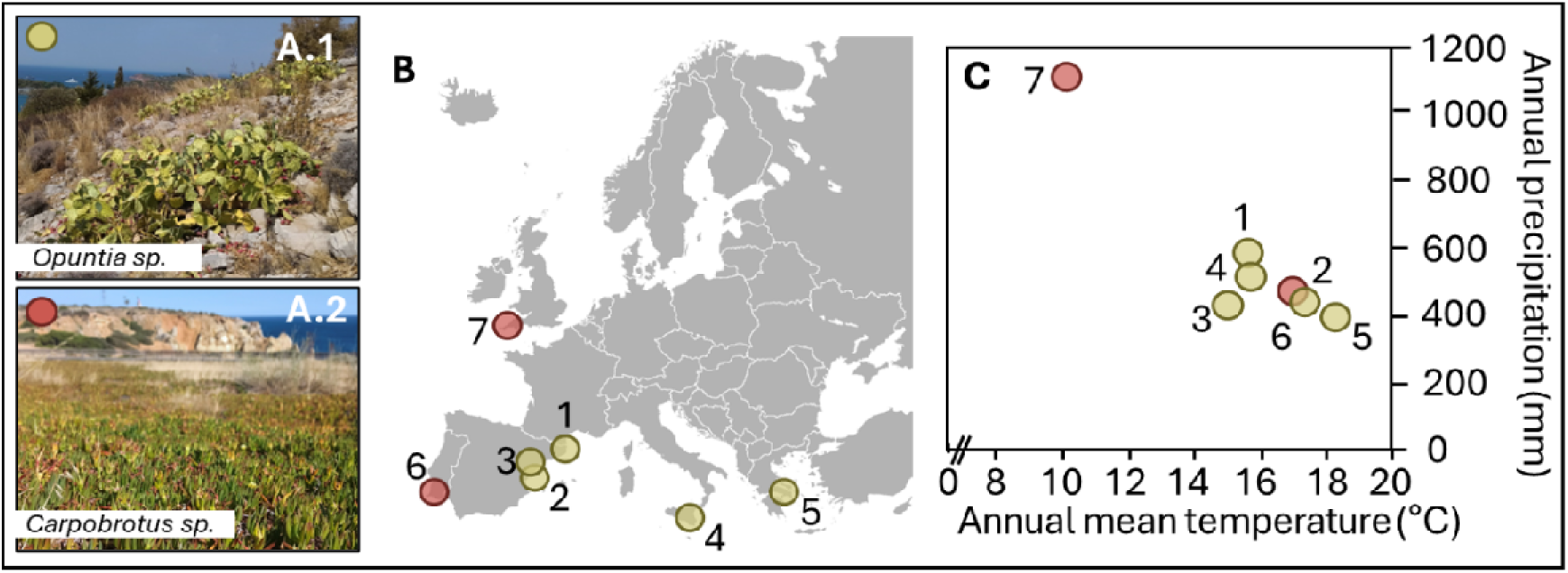
Study species (A), sites (B) and climatic conditions (C) of the studied sites. To contrast climatic conditions of the multiple study sites, we extracted annual mean temperature and annual precipitation from WordlClim (Fick and Hijmans 2017). Due to historical imagery availability, we did not use all populations for all study cases: 1: Automatic detection of the prickly pear in different habitats (Populations 1-5); 2: Spatially explicit long-term population monitoring of the prickly pear in a climatic gradient (Populations 2 and 4); and 3: Long-term individual area occupancy dynamics of the cape fig in two differentiated environments (Populations 6 and 7).

### Google Earth high resolution imagery download

To evaluate the presence of the detected individuals in previous years, we obtained public aerial imagery of the study sites from Google Earth Pro (Step 2 **Figure 1**, **Supplementary Material 1**). We selected historical Google Earth Pro imagery based on proximity to survey dates and optimal resolution (15–30 cm/pixel; **Table 1**).

### Individual segmentation

We used GPS points to segment individuals in UAV and Google Earth imagery using the GeoSAM QGIS plugin (ViT-L SAM model; Zhao et al. 2023) within QGIS 3.36.2 (QGIS.org, Switzerland). GeoSAM enables AI-assisted instance segmentation. Segmented areas were converted to bounding boxes for model training.

### Multitemporal images georeferencing and individuals matching

We georeferenced downloaded images using ≥50 nearby reference points in the GDAL Georeferencer plugin in QGIS (**Supplementary Material 1**). Thin plate spline transformation was applied to align imagery, as recommended for non-linear corrections (Bogdan et al. 2021). We then tracked individuals across years based on overlap and selected the clearest image (highest resolution and minimal occlusion) for size estimation. Individual recruitment was inferred when individuals were present in a given year but absent the previous one. Detection of small individuals is limited by resolution and occlusion and discussed further.

### Automatic detection using the AutoDL Deep learning Toolbox

Bounding boxes from the clearest Google Earth image were used to train object detection models in ArcGIS Pro (v3.4, ESRI, USA). We labelled data as PASCAL VOC and applied rotation-based augmentation. The AutoDL tool tested multiple architectures (e.g., YOLOv3, RetinaNet, Faster R-CNN) and hyperparameters to identify the best-performing model based on mean Average Precision (mAP, average precision values across all classes). We then applied the best model to detect additional individuals beyond training areas.

We tested correlations between mAP and variables including image resolution, survey area, number of individuals/images, inter-individual distances, and average bounding box size. To evaluate the visual contrast of *Opuntia* individuals, we measured hue, saturation, and brightness (HSB) in 50 plants and 50 background points using ImageJ. Unlike RGB, the HSB space aligns better with human colour perception (Kumar et al. 2016). Multivariate colour variability was analysed via multidimensional scaling using the *vegan* R package (**Supplementary Material 3**).

## 3. Results

### 3.1. Study case 1. Automatic detection of the prickly pear in different habitats

High accuracy was achieved for the automatic detection of prickly pear using public imagery from Google Earth. Across populations, the best accuracies (*i.e.* the percentage of detected individuals) ranged from 58 to 73% (**Table 1**). The architecture that resulted with the highest accuracy detecting prickly pear individuals was FoveaBox, an anchor-free object detection system that learns the object existing possibility and the bounding box coordinates without anchor reference (Kong et al. 2020).

Low colour variability of the target objects increased model accuracy (**Table 1**). Higher accuracy was observed when the colour variability of the target species was low, and the ratio of colour variability between the background and the target objects was high. Overall, variables related to the colour contrast resulted in high correlation coefficients, whereas variables such as image resolution, distance between individuals, their size and the number of training images weren’t that determinant.

**Table 1.** Deep learning for detecting invasive individuals of *Opuntia* sp. with Google Earth Imagery across multiple sites. For each site, image metadata, size of the training dataset, attributes of individual’s morphometry and colour contrast attributes between targets and background. GSD: ground sampling distance. Correlation between accuracy and each descriptor was tested using Pearson correlation. The colour scale from blue to red responds to the magnitude of the correlation coefficient.

|  |  | Opuntia sp. site |  |  |  |  | Correlation with accuracy |  |
| --- | --- | --- | --- | --- | --- | --- | --- | --- |
|  |  | 1 | 2 | 3 | 4 | 5 | Pearson coef | P-value |
| Image metadata | Image date | Oct-20 | Nov-22 | May-23 | Sep-22 | May-22 | NA | NA |
|  | GSD (cm/pixel) | 7.90 | 14.90 | 12.20 | 14.20 | 7.00 | 0.51 | 0.376 |
|  | Surveyed area (ha) | 1.27 | 1.91 | 0.84 | 0.88 | 0.82 | 0.39 | 0.513 |
| Training data | Number of individuals | 192 | 79 | 138 | 114 | 126 | -0.06 | 0.921 |
|  | Number of images | 1152 | 484 | 824 | 684 | 596 | 0.13 | 0.831 |
| Individuals' size and density | Mean individuals' distance (m) | 5.18 | 6.07 | 10.29 | 7.48 | 2.66 | 0.13 | 0.835 |
|  | Individuals > 4 m<sup>2</sup> (%) | 25.68 | 79.75 | 88.10 | 64.91 | 38.89 | -0.03 | 0.956 |
|  | Bounding box average size (m2) | 10.51 | 11.53 | 18.64 | 10.11 | 9.34 | -0.39 | 0.514 |
| Level of contrast between target individuals (OP) and background (BKG) | OP colour mean distance | 0.18 | 0.23 | 0.35 | 0.24 | 0.28 | -0.72 | 0.169 |
|  | BKG colour mean distance | 0.10 | 0.17 | 0.30 | 0.16 | 0.28 | -0.85 | 0.066 |
|  | OP vs BKG colour mean distance | 0.16 | 0.23 | 0.35 | 0.24 | 0.30 | -0.74 | 0.151 |
|  | OP colour variability | 38.42 | 48.63 | 101.25 | 52.6 | 90.27 | -0.88 | 0.050 |
|  | BKG colour variability | 62.00 | 68.61 | 124.54 | 79.27 | 82.38 | -0.59 | 0.297 |
|  | BKG/OP colour variability | 1.61 | 1.41 | 1.23 | 1.51 | 0.91 | 0.88 | 0.047 |
|  | OP-BKG colour variability | -23.58 | -19.98 | -23.29 | -26.66 | 7.88 | -0.70 | 0.186 |
| Object detection using deep learning accuracy |  | 0.687 | 0.681 | 0.602 | 0.726 | 0.584 |  |  |

The best model was obtained when training the deep learning model with the individuals identified in Sagunt (Spain), with a 73% accuracy (**Figure 3**). Using the trained model, we identified 401 individuals of *Opuntia* sp. growing near the surveyed area, which we indeed observed when we visited the area. While most detections constituted true positives (**Figure 3C**), some false positives were detected, which corresponded to rocks and shadowed areas (**Figure 3D**).

Individual size was a key determinant for successful detection. Whereas the smallest individuals (bounding box < 7 m^2^ and individual area < 3.9 m^2^, which represented <20% of the total individuals) were undetected by the best models, detection accuracy for bigger individuals (higher than the global median size) reached 80.24% (65 out of 81 individuals) (**Figure 3B**).

**Figure 3.**
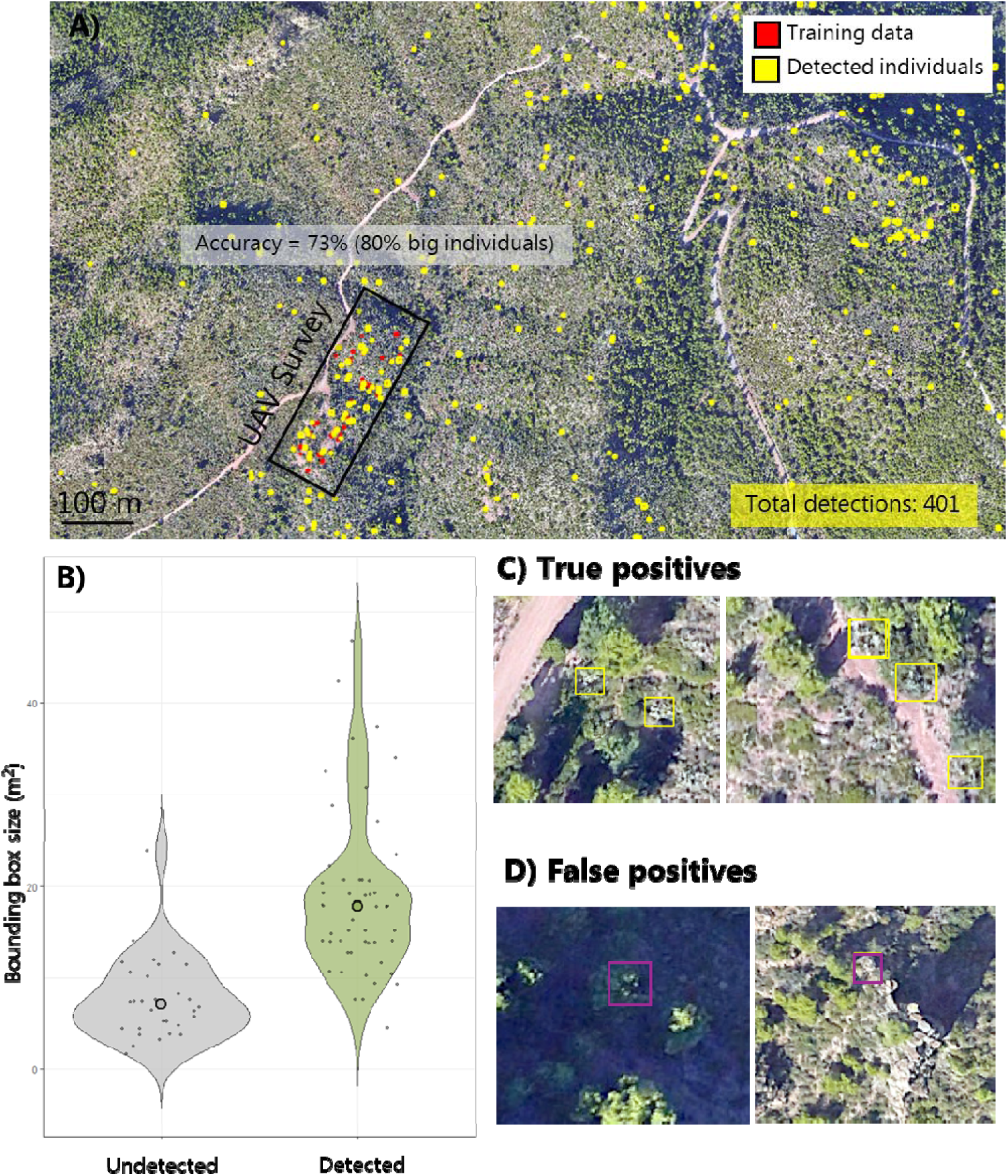
Potential of Google Earth imagery to automatically detect *Opuntia* sp. over large territories. A) Selected study area in Sagunt (Valencia, Spain) from Google Earth imagery in November 2022 (GSD = 19 cm), where the UAV survey was conducted to acquire high resolution RGB images as well as GPS positions of individuals within the black square. Red bounding boxes highlight training samples for the deep learning models, where yellow bounding boxes were the predicted result with a training accuracy of 73%. B) Violin plot of the bounding box size (pixels) of detected and undetected individuals within the training area. Big dots represent population median, revealing a clear detection bias towards bigger individuals. Accuracy of individuals bigger than the undetected median scaled to 80%. C) and D) examples of true and false positives observed outside the training area respectively.

### 3.2. Study case 2. Spatially explicit long-term population monitoring of the prickly pear in a climatic gradient

High quality Google Earth historic imagery was acquired for three prickly pear populations along a climatic gradient, obtaining clear images with at least 30 cm/pixel since 2012, in most cases with annual frequency. Individual’s matching from 2024 observations led to identify between 79 and 126 individuals in the best Google Earth image. The best images were obtained for the population in Greece, where from the 114 individuals detected in 2022, only 50 already existed in 2014 (**Supplementary material 2**).

Spatially explicit individual tracking across time led to a set of potential applications for ecological and conservation studies including the ability to estimate individual age, track population growth, identify high and low productive years (in terms of recruitment), characterise population structure, model age-size relationships, and identify high and low recruitment areas (**Figure 4**).

Annual recruitment rates ranged from 1.4 to 8.5 individuals/year across the three populations studied (**Figure 4E**). Aridity correlated strongly with individual recruitment (P-value < 0.01, R^2^ > 0.88), revealing how the population with higher mean annual temperature and lowest annual precipitation showed the strongest recruitment rates (**Figure 4E**, Supplementary Material 4).

**Figure 4.**
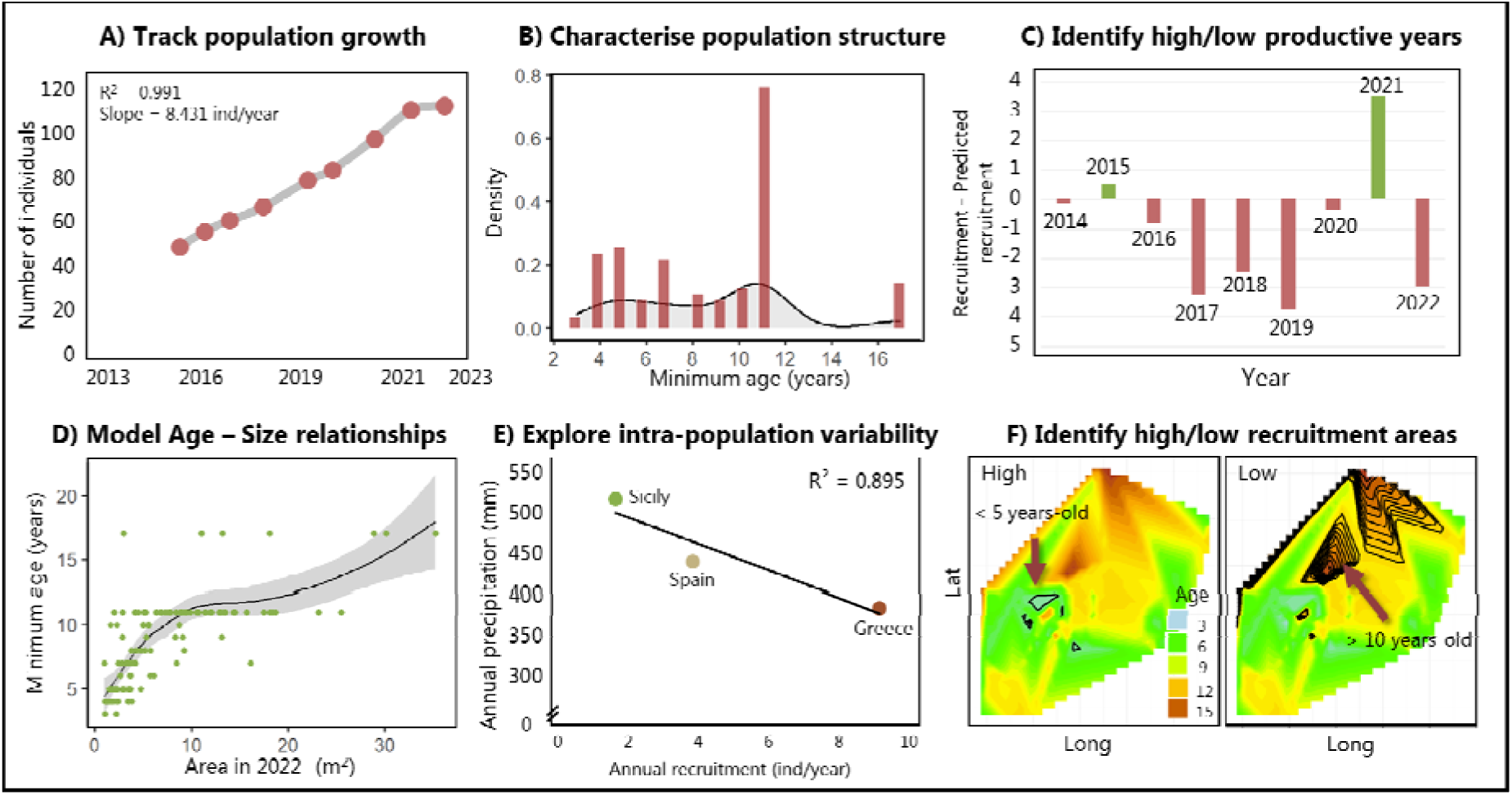
Applications of retrospective image analysis for invasive species management. Example of the applications on an invasive cacti population in Greece, using 10-years imagery (2024-2014). A) Track population growth: by checking species presence back in time we can acquire recruitment data. B) Characterise population structure: by acquiring sizes of the individuals, we can understand population structure. C) Identify high and low productive years: by contrasting predicted versus observed recruitment we can identify productive years and fit models to test their dependence on climatic or other variables. D) Model the relationship between age and size. By tracking individuals back in time, we can assign age and model size increase to predict further individual age. E) Recruitment of prickly pear in three populations with a climatic gradient (from more arid, Greece to more humid Sicily). F) Since recruitment data is geolocalised we can map recruitment and detect high or low recruitment areas to prioritize managing.

### 3.3. Study case 3. Long-term individual area occupancy dynamics of the cape fig in two differentiated environments

Quantification of invaded area to plan management costs is possible with Google Earth imagery. In 2025, a total area of 7,350.42 m^2^ of *Carpobrotus* sp. was observed in Ponta da Piedade (Portugal), and 639.27 m^2^ in Gwynver Beach (UK) in 2022, corresponding to the most up to date high quality Google Earth image (**Supplementary Material 5**). According to a recent estimation of the mean biomass per area of this species (Figuereido Meyer et al. 2023), those areas might translate into 71.15 and 6.19 t of fresh weight, respectively (**Figure 5**).

Invaded area expansion or shrinkage can also be monitored from Google Earth imagery (**Figure 5**). The cape fig showed an overall increase in area in the two contrasted environments, revealing the impressive amount of 18 m^2^ expansion/year in the coldest region (from 2017-2022). Since the species show strong asexual reproduction creating dense and large mats, genet (a collection of physiologically independent units sharing a genotype) monitoring was only possible for determined patches that grew in undisturbed habitats and displayed clear limits (**Figure 5**). From a set of 15 individuals, we analysed area occupancy dynamics, revealing a consistent mean growth rate of 12.95 ± 5.32 m^2^/year (**Figure 5F**). Whereas smaller patches showed smaller and positive growth rates, higher variability in transient area changes were observed for bigger genets (**Figure 5D, E**).

**Figure 5.**
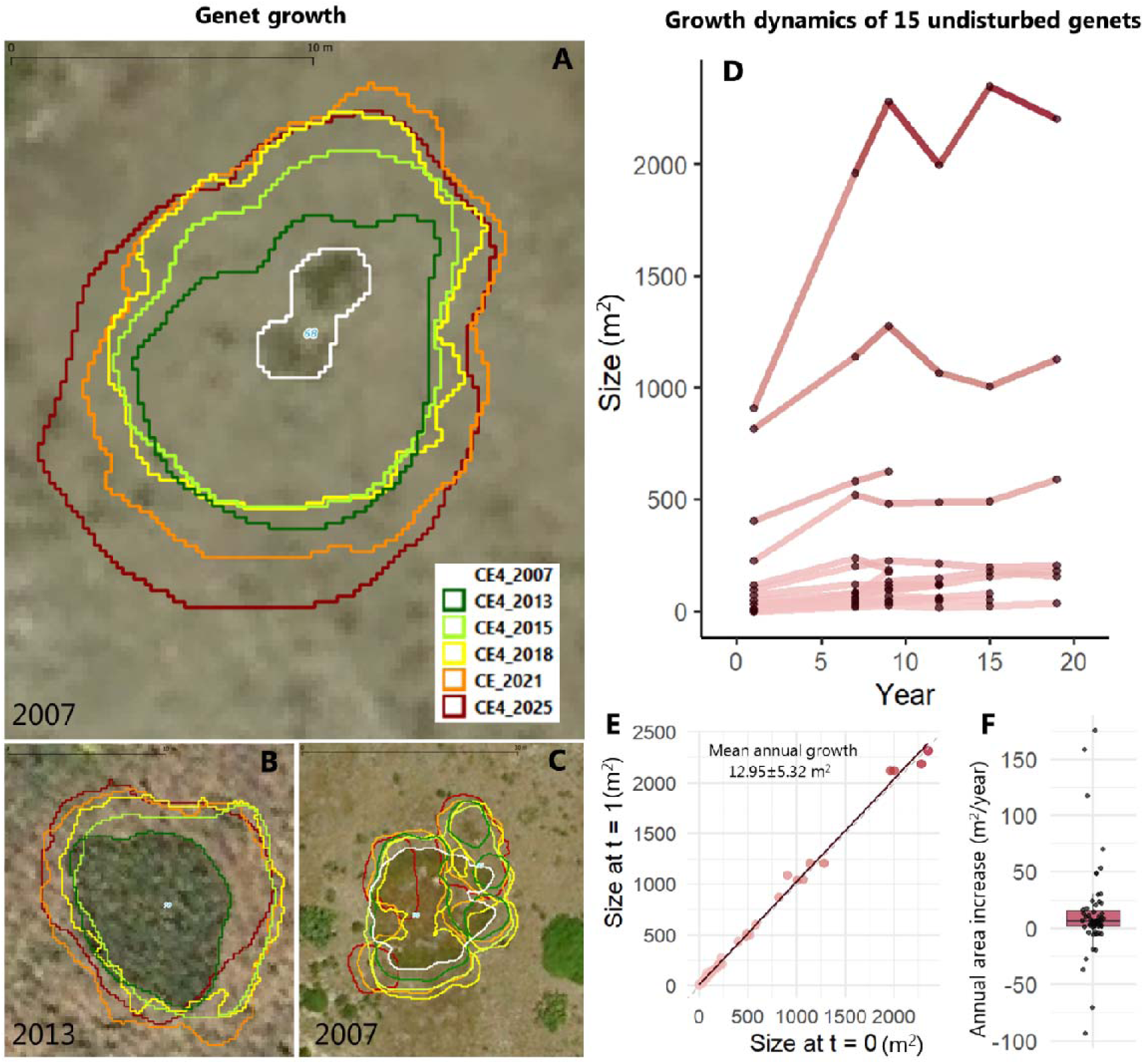
Genet growth dynamics in Site 1 (Ponta da Piedade, Portugal). A-C examples of segmented sizes across years. Individuals A and C were already present in 2007, whereas B was first registered in 2013. Some genets decreased in their area due to fragmentation (as C), whereas others were not found due to disturbance. D) Area changes in time for all the 15 monitored genets for almost 20 years. E) Growth transitions from a given time to the next transformed to annual rates coloured by size. The grey line represents size maintenance 1:1. The black line is a fitted linear model (R^2^ = 0.995). F) Annual area changes boxplot, showing mean 12.95 ± 5.32 m^2^/year across the 15 individuals on all time steps.

## 4. Discussion

Long-term population monitoring remains challenging due to logistical constraints, high costs, and limited data availability. Here, we introduce an analytical pipeline to collect retrospective, long-term demographic data using publicly available Google Earth imagery and deep learning. We demonstrate its potential by tracking individuals of two invasive plant species with contrasting growth forms across 20 years, revealing sustained recruitment and climate-driven responses. This pipeline provides a low-cost, scalable tool for monitoring biological invasions and ecological change.

Limited training data is one of the challenges for successfully training deep learning algorithms for object detection (Pichler and Hartig 2023). The quality and quantity of high-resolution public aerial imagery now allow for a significant expansion in the number of images of a specific species. Invasive species tend to rapidly colonize an area generating high density populations (Arim et al. 2005). Therefore, plenty of individual images can be obtained from a relatively small area (Rakgoale and Ngetar 2024), as we demonstrated with *Opuntia* sp. and *Carpobrotus* sp. in multiple sites across Europe, where we tracked 319 prickly pear individuals and *ca*. 8,000 m^2^ of *Carpobrotus* sp.

Effective population monitoring requires long-term efforts (Blanc and Thrall, 2024; Stroud and Ratcliff, 2025), often exceeding the duration of funding cycles (Lindenmayer et al. 2012). Typical demographic studies would start at the present time and collect data for multiple time intervals (Gamelon et al. 2021). Imagery-based demography using drones has gained popularity in the last decade (Gomes et al. 2018; Bogdan et al. 2021; Hao et al. 2024; Olsoy et al. 2024), but it is restricted to multitemporal site visits and permission approvals. Generating long-term demographic series that ensure meaningful trends thus takes at least 10 years of sustained funding and expertise. However, our methodology provides a tool to acquire demographic data differently, by traveling back in time.

The high resolution of Google Earth imagery improves detection accuracy and allows identifying smaller species compared to satellite-based detection, while significantly reducing costs. Unlike satellite imagery, which often uses pixel-based approaches that cannot delineate individuals (Kattenborn et al. 2020), Google Earth images support individual-based analysis. For instance, Lake et al. (2022) used Worldview-2 (0.5 m) and Planetscope (3 m) imagery to detect areas with *Euphorbia virgata* but could not resolve individuals. Deep learning approaches applied to satellite data often rely on multispectral bands (e.g. Duncan et al. 2023; Zagajewski et al. 2024), whereas species detection from Google Earth requires distinct RGB or textural features (Müllerová et al. 2017). While its resolution enables both pixel- and object-based detection, Google Earth imagery does not replace high-resolution multispectral satellites (Sankey et al. 2017) but offers a cost-effective alternative to targeted RGB drone surveys (e.g. Müllerová et al. 2017; Elkind et al. 2018; Amarasingam et al. 2024; Gautam et al. 2025), with broad spatial coverage (Minnich et al. 2011; Watanabe et al. 2020).

Species detectability depends on phenology and image timing. For example, *Opuntia* species are more visible in summer when surrounding vegetation is dry (Müllerová et al. 2017; Marzialetti et al. 2021). Oddi et al. (2021) showed that detection accuracy dropped significantly between August and October, highlighting the importance of seasonal image selection. Our method—combining UAV-based ground-truth segmentation with historical Google Earth imagery—offers a cost-effective balance between spatial coverage and resolution for individual-level monitoring.

However, image quality and resolution may vary across time, particularly with opportunistic or publicly available imagery, limiting the detection of small individuals, seedlings, or subtle phenological changes. Shadows, occlusions, and misalignments between time points can further reduce accuracy. Without field validation, confirming individual presence or absence is difficult, meaning mortality events may go undetected, as only survivors are traced backwards. Retrospective image analysis is therefore most effective when paired with ground-truth data (Gränzig et al. 2021) or applied to species that are large, sessile, spatially distinct, and have visually distinct traits (Müllerová et al. 2017). In some cases, full demographic models may be feasible depending on species traits and image resolution. For example, *Carpobrotus* populations were analysed using 0.35 cm/pixel imagery across two consecutive years to build an integral projection model (IPM), enabling estimates of growth, survival, reproduction, and flower counts (Bogdan et al. 2021).

Retrospective image analysis offers a non-invasive approach to extract key demographic parameters (Visser et al. 2014; Munteanu et al. 2024). As shown with *Opuntia* sp., this approach can yield estimates of individual’s age, recruitment over time, changes in population size and spatial distribution as well as mortality events. For *Opuntia* sp. we showed consistent recruitment rates across years, suggesting population growth especially in arid areas, where recruitment rates escalated to 8.5 individuals/year. Our methodology allowed us to infer individual age or at least minimum age on species whose age cannot be estimated through dendroecological lens, revealing a size-age model that could be used to infer approximate age of further individuals (Cao et al. 2018) allowing further ecological and evolutionary analysis on the universality of senescence (Munné-Bosch, 2015). When image resolution allows, it is also possible to quantify individual growth rates and detect life stage transitions, especially in size-structured populations. For clonal species, such as *Carpobrotus* sp., retrospective analysis can reveal patterns of clonal expansion or contraction, ramet persistence, and fragmentation. Moreover, long-term image series can uncover responses to disturbances, environmental shifts, or management interventions, offering a powerful lens into population resilience and trajectory without requiring continuous field presence.

Spatiotemporal population analysis provides a critical framework for understanding ecological and evolutionary processes across landscapes and through time. By integrating spatial and temporal dimensions, our pipeline can quantify how population structure, density, and dynamics vary in response to environmental heterogeneity. One application of spatiotemporal data is mapping vital rates (*e.g*., survival, growth, reproduction) across different scales. For example, Cao et al. (2018) built a map of survival and regeneration of *Caragana* sp. in central Inner Mongolia based on imagery-based demography on multiple populations along a precipitation and temperature gradient. Similarly, in our study we report aridity-dependant recruitment in *Opuntia* sp. when analysing large spatial scale and identified low and high recruitment areas at the local spatial scale, providing essential information to fit mechanistic distribution models (Fenollosa et al. 2025b; Briscoe et al. 2019) and establishing management priorities. Further applications of our proposed methodology go beyond invasion science (Visser et al. 2014) and include remote areas or rare species surveying (Rominger et al. 2021; Rominger and Meyer, 2021), monitor ecosystem restauration (Robinson et al. 2022; Singh et al. 2024), characterise management effects at the community level (Pasquarella et al. 2016; Jaime et al. 2023) or early-warning signals for biodiversity management (Alibakhshi, 2023).

Retrospective image analysis for spatiotemporal modelling may not be suitable for all species, habitats and regions, but can be easily implemented for some. Species characteristics together with habitat and region will determine applicability (**Figure 6**). Key species attributes include large area, distinctive colour and texture from the surroundings, sessile, perennial, with high population growth rates, low turnover, and high abundance. Open, sparsely vegetated habitats such as dunes, arid or semiarid biomes, tundra or high mountain constitute ideal scenarios (Mao et al. 2021). In **Figure 6**, we provide a list of already identified invasive plant candidates that would meet all criteria. At the regional scale, Google Earth high resolution and multitemporal imagery are not equally available around the globe, but some alternatives can complement aerial data (Lesiv et al. 2018). Other sources of high-resolution muti-temporal imagery are Google Street View imagery (Li 2020), Microsoft Bing Maps (Lesiv et al. 2018), GeoNadir (https://data.geonadir.com/) or even spy-satellite images (Munteanu et al. 2024).

The proposed methodology is not restricted to big plant taxa. In general, species that have been successfully detected from satellite imagery may be suited for retrospective image analysis from Google Earth imagery. Beyond species, our pipeline can be used to monitor population size of some moving species, such as grey seals (Moxley et al. 2017) and use inverse estimation to build demographic models from species population-level data (González et al. 2016). However, in those cases, we would need higher resolution on all years, instead of relying on ground truth detection on the present moment and trace back individuals. Indirect elements can be used as well to track back population dynamics, such as seabirds’ nests (Lalach et al. 2023), insect-infected trees (Cardil et al. 2017), marmot burrows (Koshkina et al. 2019), beaver dams and lodges (Graham et al. 2022), and guano stains (Witharana and Lynch, 2016).

**Figure 6.**
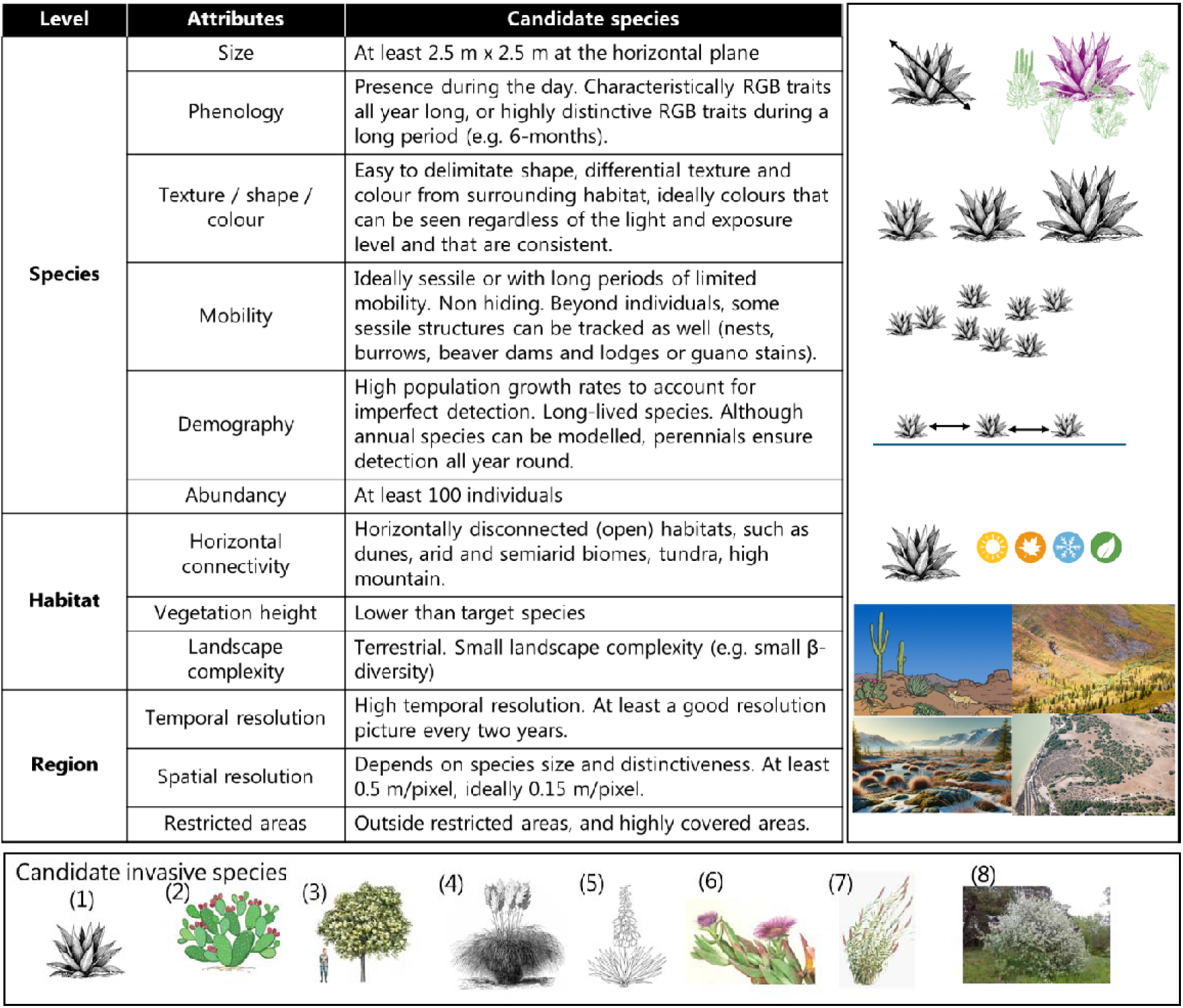
Species, habitat and region requirements to acquire demographic data from retrospective image analysis. Bottom panel: List of eight invasive species that constitute ideal candidates: (1) *Agave americana*, (2) *Opuntia stricta*, *Opuntia ficus-indica*, *Opuntia* sp. (3) *Pinus radiata*, (4) *Cortaderia selloana,* (5) *Yucca* sp., (6) *Carpobrotus edulis*, *Carpobrotus acinaciformis*, (7) *Arundo donax*, (8) *Elaeagnus angustifolia*.

## Acknowledgments

We thank C. Ribalta-Pizarro for her assistance geolocalising individuals on the field and collecting UAV data. This project is funded by UKRI Marie Skłodowska-Curie Actions Fellowship (EP/Y02873/1) of E.F. hosted by R.S.G. R.S.G. was supported by a NERC Pushing the Frontiers grant (NE/X013766/1).

## 5. Data Availability

Data will be published in FigShare. Insert DOI here.

