## Supplementary material 1-5 for "Retrospective image analysis for long-term demography using Google Earth imagery"

**Supplementary material 1**: Protocol and set of videos on how to perform the analysis. ZIP folder. Video-tutorials can also be found in this YouTube playlist: https://www.youtube.com/playlist?list=PL_LKE-yTi9kBXfw_qDdJCQ3Sxu2fjGvDD, of EF’s account: @environmentaldatascientist.

**Supplementary material 2:** Example of individuals tracking from public imagery in a population of *Opuntia* sp. in Greece (Population 4). Example of individuals across time.

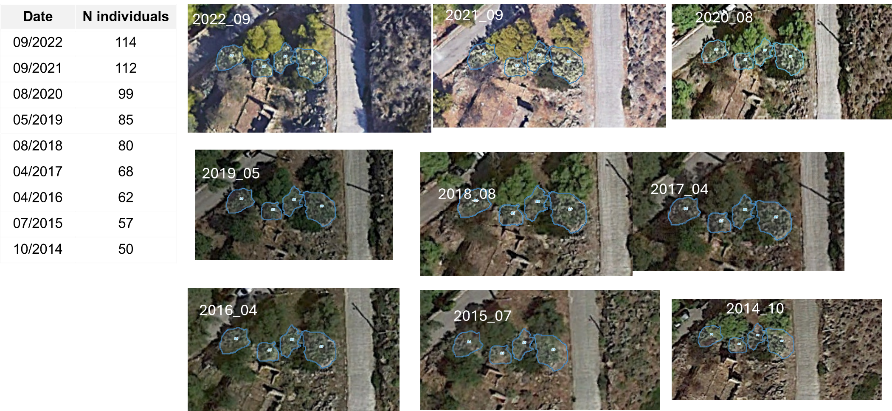

**Supplementary material 3**: Multidimensional analysis of target object colour contrast. In order to quantify colour contrast between the target objects (*Opuntia* sp. individuals) and the surroundings we extracted colour coordinates (mean and median) within the HSB colour space of 50 random background points and 50 *Opuntia* sp. individuals at each studied location. Colour features were used to perform multidimensional scaling and quantify the level of multivariate colour variation within and between groups (sites * Background/Target). The MDS plot shows the dissimilarity between targets (reddish colours) and background points (greenish colours) in each site.

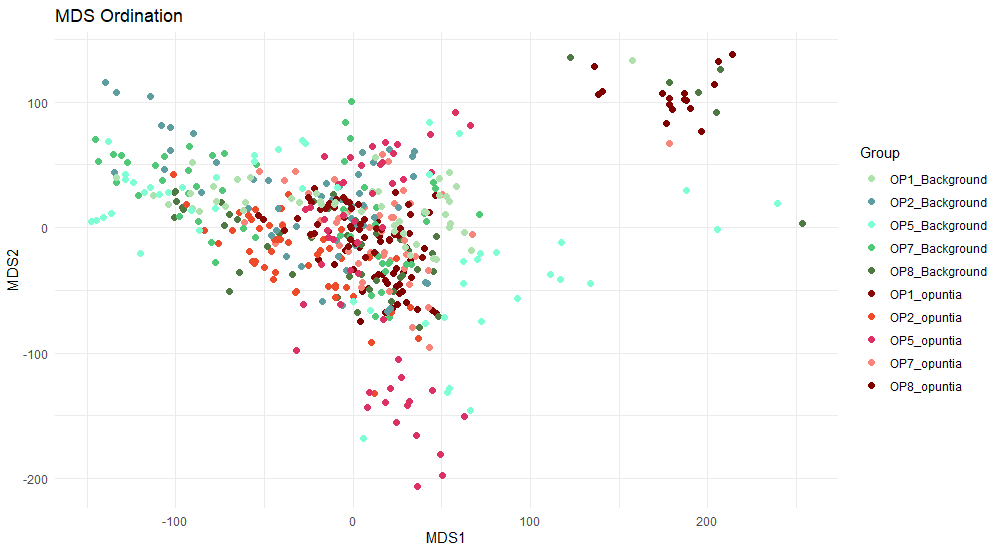

**Supplementary material 4**

Recruitment of prickly pear in three populations with a climatic gradient. Populations are coloured by aridity (from more arid, Greece to more humid Sicily). A) Number of individuals tracked across years. B and C) annual recruitment rates correlation with annual mean temperature and annual precipitation.

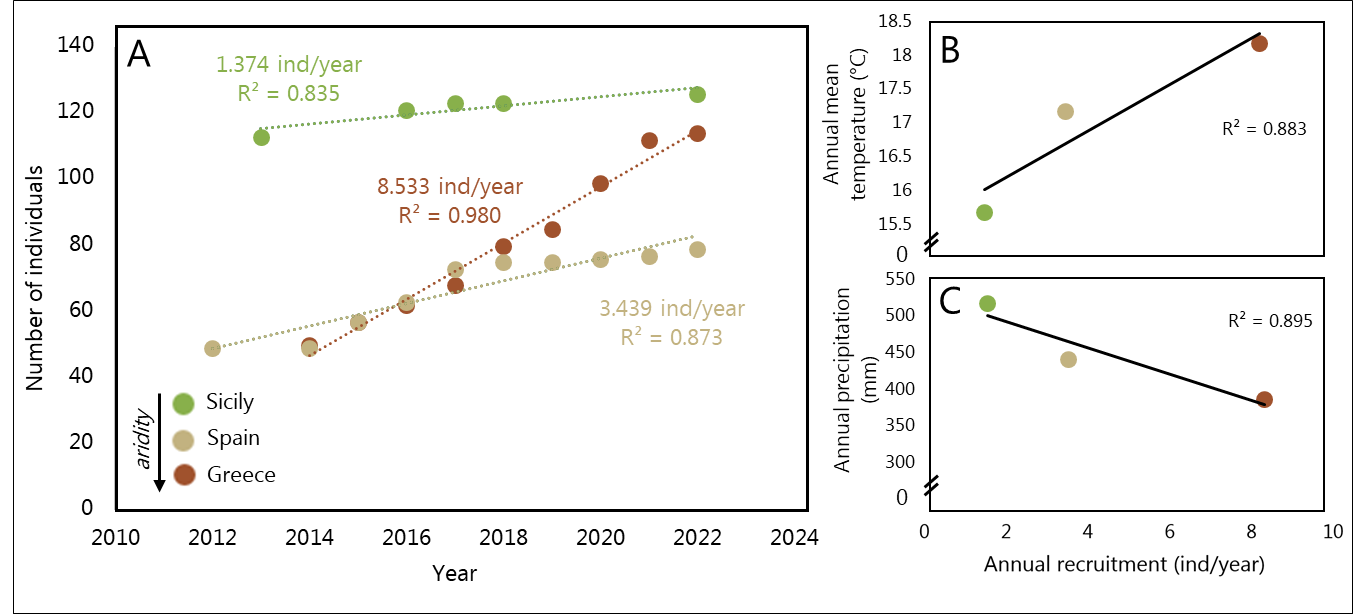

**Supplementary Material 5**

Detected *Carpobrotus* sp. area across two populations in Portugal and the UK. The upper images area best resolution Google Earth imagery of the area, whereas bottom images are drone images taken in summer 2024. Letters symbolise the 15 undisturbed individuals analysed in the manuscript. Coloured lines align to segmented areas from Google Earth imagery across the multiple years.

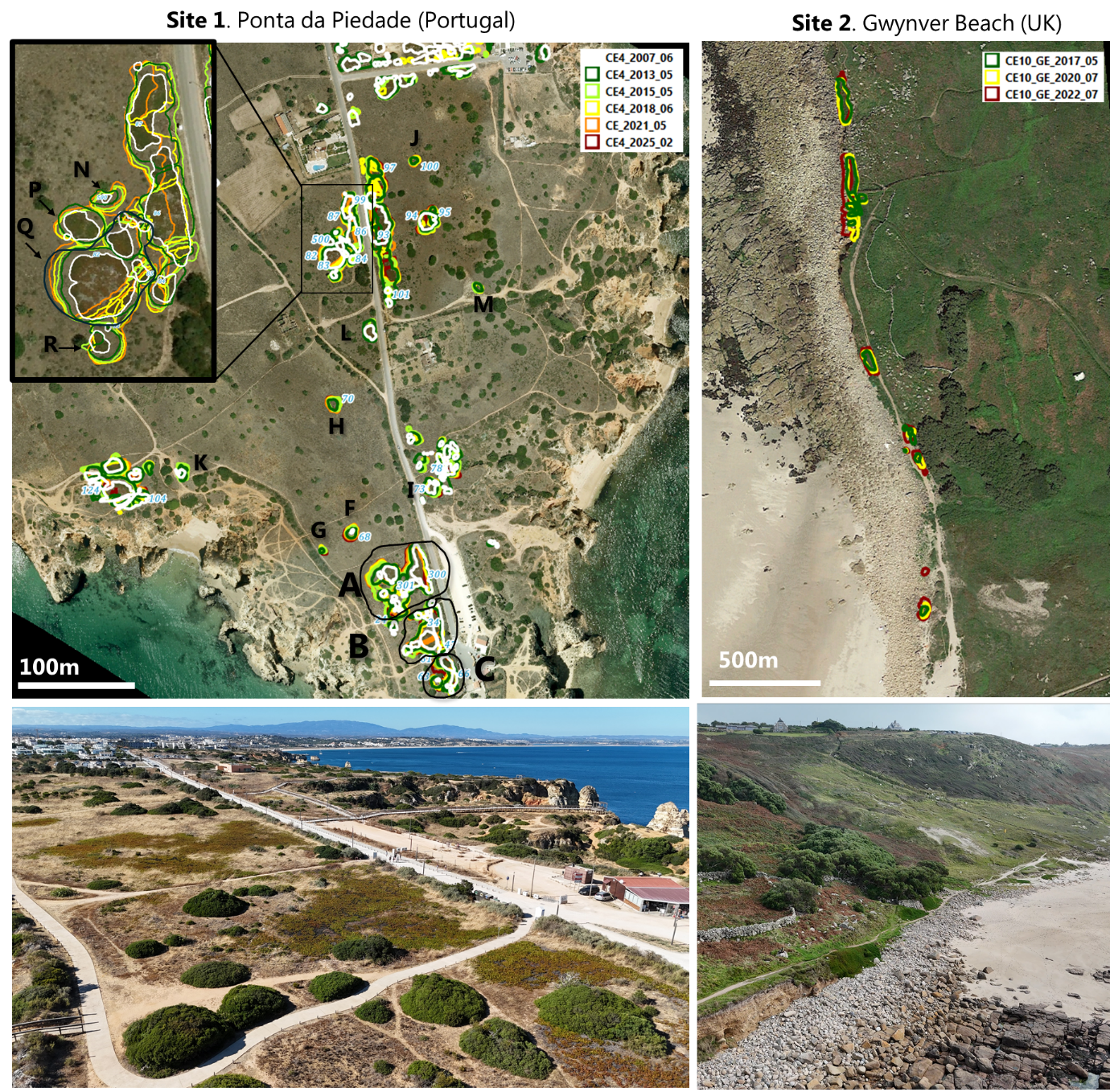
